# Forging new antibiotic combinations under iron-limiting conditions

**DOI:** 10.1101/775445

**Authors:** Derek C. K. Chan, Irene Guo, Lori L. Burrows

**Author notes:** **Correspondence to:** Dr. Lori L. Burrows, 2238 MDCL, 1280 Main St. West, Hamilton, ON L8S 4L8 Canada.

## Abstract

*Pseudomonas aeruginosa* is a multidrug-resistant nosocomial pathogen. We showed previously that thiostrepton (TS), a gram-positive thiopeptide antibiotic, was imported via pyoverdine receptors and synergized with iron chelator deferasirox (DSX) to inhibit the growth of *P. aeruginosa* and *Acinetobacter baumannii* clinical isolates. A small number of *P. aeruginosa* and *A. baumannii* isolates were resistant to the combination, prompting us to search for other compounds that could synergize with TS against those strains. From literature surveys we selected 14 compounds reported to have iron-chelating activity, plus one iron analogue, and tested them for synergy with TS. Doxycycline (DOXY), ciclopirox olamine (CO), tropolone (TRO), clioquinol (CLI), and gallium nitrate (GN) synergized with TS. Individual compounds were bacteriostatic but the combinations were bactericidal. Our spectrophotometric data and chrome azurol S agar assay confirmed that the chelators potentate TS activity through iron sequestration rather than through their innate antimicrobial activities. A triple combination of TS + DSX + DOXY had the most potent activity against *P. aeruginosa* and *A. baumannii* isolates. One *P. aeruginosa* clinical isolate was resistant to the triple combination, but susceptible to a triple combination containing higher concentrations of CLI, CO, or DOXY. All *A. baumannii* isolates were susceptible to the triple combinations. Our data reveal a diverse set of compounds with dual activity as antibacterial agents and TS adjuvants, allowing combinations to be tailored for resistant clinical isolates.

## INTRODUCTION

Iron is a critical micronutrient for bacteria, influencing biofilm formation, pathogenicity, and growth (1, 2)., The opportunistic Gram-negative pathogen *Pseudomonas aeruginosa* has an extensive repertoire of iron acquisition systems that are upregulated during iron-deplete conditions, similar to those encountered during infection. To overcome iron limitation, *P. aeruginosa* produces the iron-scavenging siderophores pyochelin and pyoverdine that bind iron with low and high affinity, respectively (3–5). Pyoverdine and its outer membrane receptors, FpvA and FpvB, are highly expressed in low-iron conditions (3, 6, 7). Pyoverdine has such a high binding affinity (10^32^ M^−1^) for iron that it can strip it from transferrin, a mammalian protein responsible for sequestering iron to impede bacterial growth (8–11). RNA-seq data showed that pyoverdine biosynthetic enzymes and uptake are highly upregulated *in vivo* in response to the iron-deprived environment (7). *P. aeruginosa* deficient in iron-uptake mechanisms are less able to cause infections compared to their wild-type counterparts (12).

Natural products often exploit iron acquisition pathways to cross the Gram-negative outer membrane. Pyocin S2, produced by *P. aeruginosa* to kill competing strains, and related toxins are taken up via FpvA (13, 14). The sideromycins, which resemble siderophores but have intrinsic antibacterial activity, also exploit iron uptake pathways (15). Taking advantage of this phenomenon, several groups have created synthetic siderophore-beta-lactam conjugates to target Gram-negative bacteria, using the iron-binding group as a Trojan horse to deliver antibiotics (16–18). One such example is cefiderocol, a siderophore beta-lactam that recently completed Phase III clinical trials. The catechol group of cefiderocol binds iron and the complex is taken up via PiuA, an outer-membrane receptor for iron transport (18, 19). The compound demonstrated potent activity against *Escherichia coli* and *Klebsiella pneumoniae* (19, 21). Thus, the Trojan horse approach enhances the delivery of antibiotics compared to diffusion alone.

Our group recently discovered that the thiopeptide antibiotic thiostrepton (TS) hijacks pyoverdine receptors under iron-limited conditions to cross the outer membranes of the World Health Organization’s top two critical priority pathogens, *P. aeruginosa* and *Acinetobacter baumannii* (22). TS activity was potentiated in heat-inactivated mouse and human serum, and by FDA-approved iron chelators, deferiprone (DFP) and deferasirox (DSX). However, a small number of *P. aeruginosa* and *A. baumannii* strains were resistant to TS-chelator combinations, prompting us to look for new compounds that could synergize with TS to inhibit those clinical isolates.

With the aim of finding compounds that could synergize with TS in iron-limiting conditions, we performed a literature search to identify bioactive iron chelators. We selected 14 putative iron-binding compounds as well as gallium, an iron analogue. Five compounds synergized with TS and had activity against *P. aeruginosa* and *A. baumannii* clinical isolates. Each compound was bacteriostatic against *P. aeruginosa* PA14; however, the addition of TS made the combination bactericidal. Growth of one highly-resistant *P. aeruginosa* clinical isolate was inhibited with higher concentrations of three of the compounds in combination with TS+DSX. These data identify a set of molecules of diverse structure and biological activity that synergize with TS, providing the ability to tailor combinations for resistant strains.

## RESULTS

### Iron-binding antibiotics form coloured complexes

We first screened a panel of common antibiotics for potential iron-chelating activity using a qualitative assay, monitoring change in colour upon addition of FeCl_3_. Binding of transition metals results in formation of coloured complexes that absorb in the visible wavelengths of light, detectable by spectroscopy and by eye (23–25). The panel consisted of 22 antibiotics from the aminoglycoside, fluoroquinolone, beta-lactam, and tetracycline classes (Fig. 1). Iron chelators DFP and DSX served as positive controls, turning dark red/violet upon addition of ferric iron at a final concentration of 10 µM. The tetracyclines – doxycycline (DOXY), tetracycline, and minocycline – exhibited similar colour changes. The fluoroquinolones – ciprofloxacin, ofloxacin, and pipemedic acid – formed orange complexes; however, the intensity of the colour change was weaker compared to the tetracyclines, DFP, and DSX. A number of beta-lactams showed colour changes ranging from a brown-orange to red-orange. Ceftriaxone was the only beta-lactam that turned red in the presence of ferric iron. Trimethoprim turned golden-yellow.

**Figure 1.**
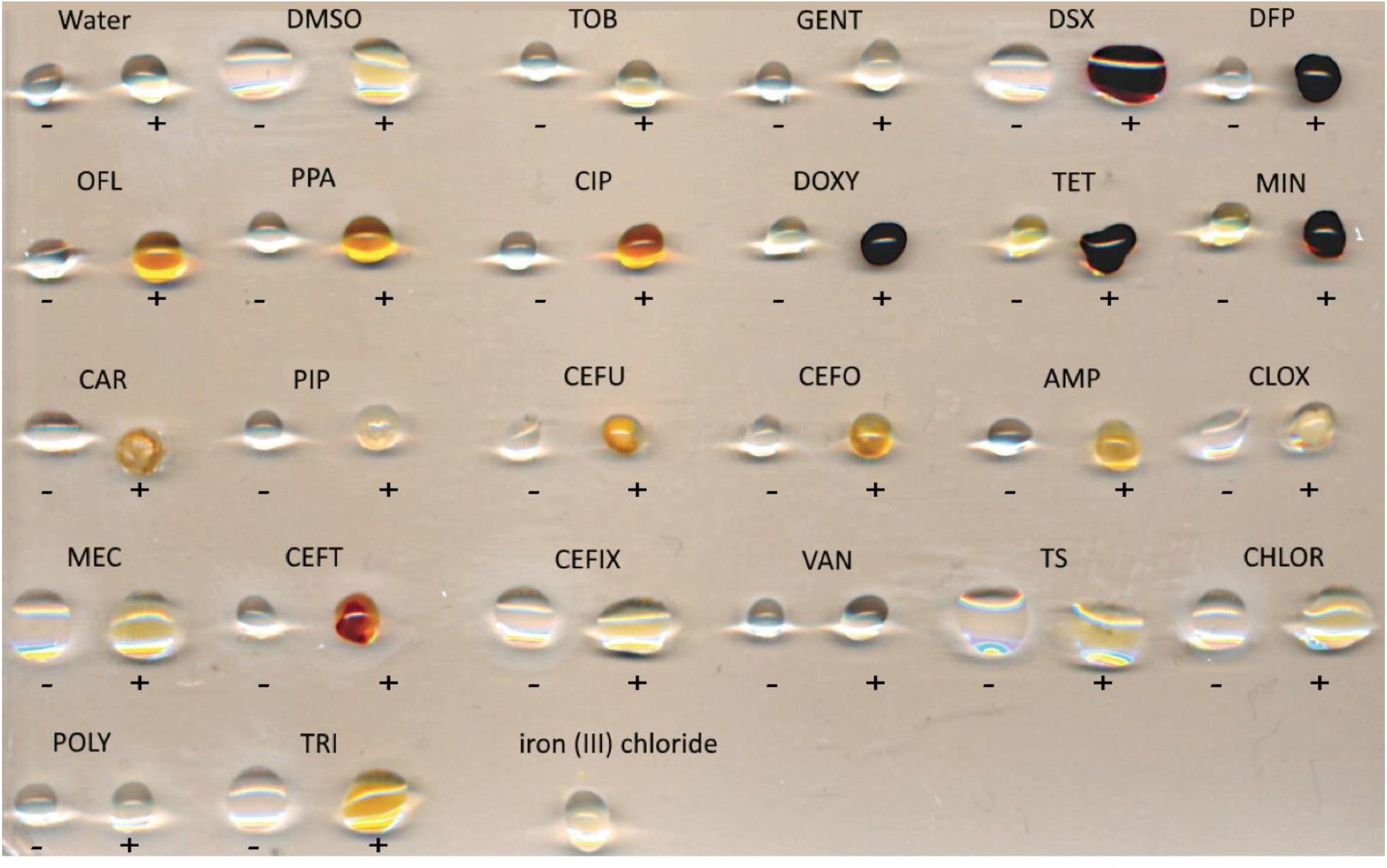
Qualitative assay to identify potential antibiotic-Fe^3+^ complexes. Binding of iron by a compound causes spectral shifts that can detected visually. Five µL of stock concentration antibiotic (below) was added to 5µL of FeCl_3_ to a final FeCl_3_ concentration of 10µM and incubated at room temperature for one hour. Negative controls without iron are indicated by a negative sign and droplets with FeCl_3_ are indicated with a positive sign. Vehicle controls with Milli-Q H_2_O and DMSO were included. The concentrations of each antibiotic stock were: TOB (tobramycin 4 mg/mL), GENT (gentamicin 10 mg/mL), DSX (deferasirox 20 mg/mL), DFP (deferiprone 60 mg/mL), OFL (ofloxacin 4 mg/mL), PIP (pipemedic acid 64 mg/mL), CIP (ciprofloxacin 5 mg/mL), DOXY (doxycycline 50 mg/mL), TET (tetracycline 20 mg/mL), MIN (minocycline 20 mg/mL), CAR (carbenicillin 100 mg/mL), PIPER (piperacillin 6 mg/mL), CEFU (cefuroxime 30 mg/mL), CEFO (cefotaxime 30 mg/mL), AMP (ampicillin 30 mg/mL), CLOX (cloxacillin 30 mg/mL), MEC (mecillinam 30 mg/mL), CEFT (ceftriaxone 30 mg/mL), CEFIX (cefixime 12 mg/mL), VAN (vancomycin 30 mg/mL), TS (thiostrepton 20 mg/mL), CHLOR (chloramphenicol 50 mg/mL), POLY (polymyxin B 4 mg/mL), TRI (trimethoprim 50 mg/mL).

### Binding of ferric iron shifts absorption spectra

To verify spectral shifts for compounds that changed colour upon addition of ferric iron, a 96-well spectrophotometric assay was performed, with final concentrations of antibiotic and FeCl_3_ of 300 µM each. The absorption spectra were scanned from 300 – 700 nm. The spectra of ciprofloxacin (CIP), pipemedic acid, ofloxacin, tetracycline, minocycline, DOXY, DSX and DFP shifted after the addition of FeCl_3_ (Fig. 2), confirming the results of the qualitative assay. Chloramphenicol and ampicillin served as negative controls. The spectrum for ceftriaxone did not change at the concentrations tested, suggesting that the changes in color observed for beta-lactams were likely due to concentration effects.

**Figure 2.**
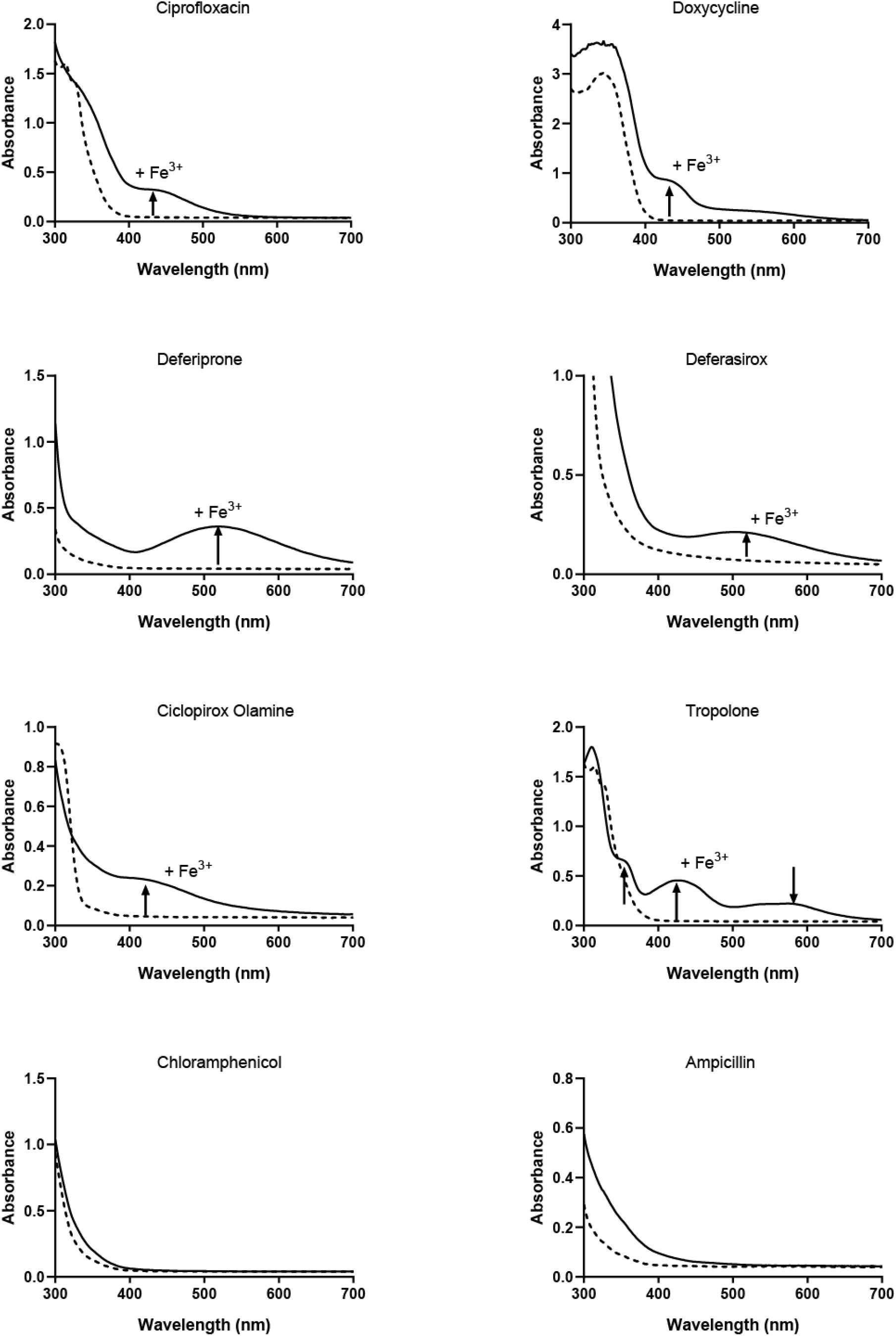
UV-Vis absorption spectrum of compounds with and without Fe (III). Equimolar concentrations of compound and FeCl_3_ were added in deionized H_2_O to a final concentration of 300 µM and a spectrum of wavelengths from 300 nm to 700 nm read after 1 h incubation at room temperature. The black dashed line is the spectrum of the compound in the absence of iron. The black solid line is the spectrum after the addition of iron. New peaks appearing after the addition of Fe^3+^ are indicated with arrows. Chloramphenicol and ampicillin were used as negative controls. Each assay was performed at least 3 times and averaged values are shown.

### Identification of other compounds that chelate iron

To expand our panel of potential chelators beyond known antibiotics, we searched the literature for bioactive compounds that were reported to have iron-chelating activity. We identified 14 compounds (Table 1) plus gallium nitrate (GN). Gallium is an iron analogue that inhibits siderophore production, iron uptake, and the activity of enzymes that use iron (26). The spectrophotometric assay was repeated for all compounds listed in Table 1 except for clioquinol (CLI). CLI was identified in our previous screen as a *P. aeruginosa* growth inhibitor (22) but precipitated at concentrations above 8 µg/mL. Ciclopirox olamine (CO) and tropolone (TRO) showed shifts in their absorption spectra (Fig. 2). A chrome azurol S (CAS) assay was also used to detect iron binding through de-colourization of the blue agar, indicating removal of Fe^3+^ from the CAS-HDTMA complex (**Fig. S1**). DSX, TRO, and CO showed the greatest decolourization, and thus the highest relative affinity for iron. Interestingly, DOXY showed a marked colour shift in the presence of Fe^3+^ (Fig. 2) but minimally decolourized CAS agar.

**Table 1:**
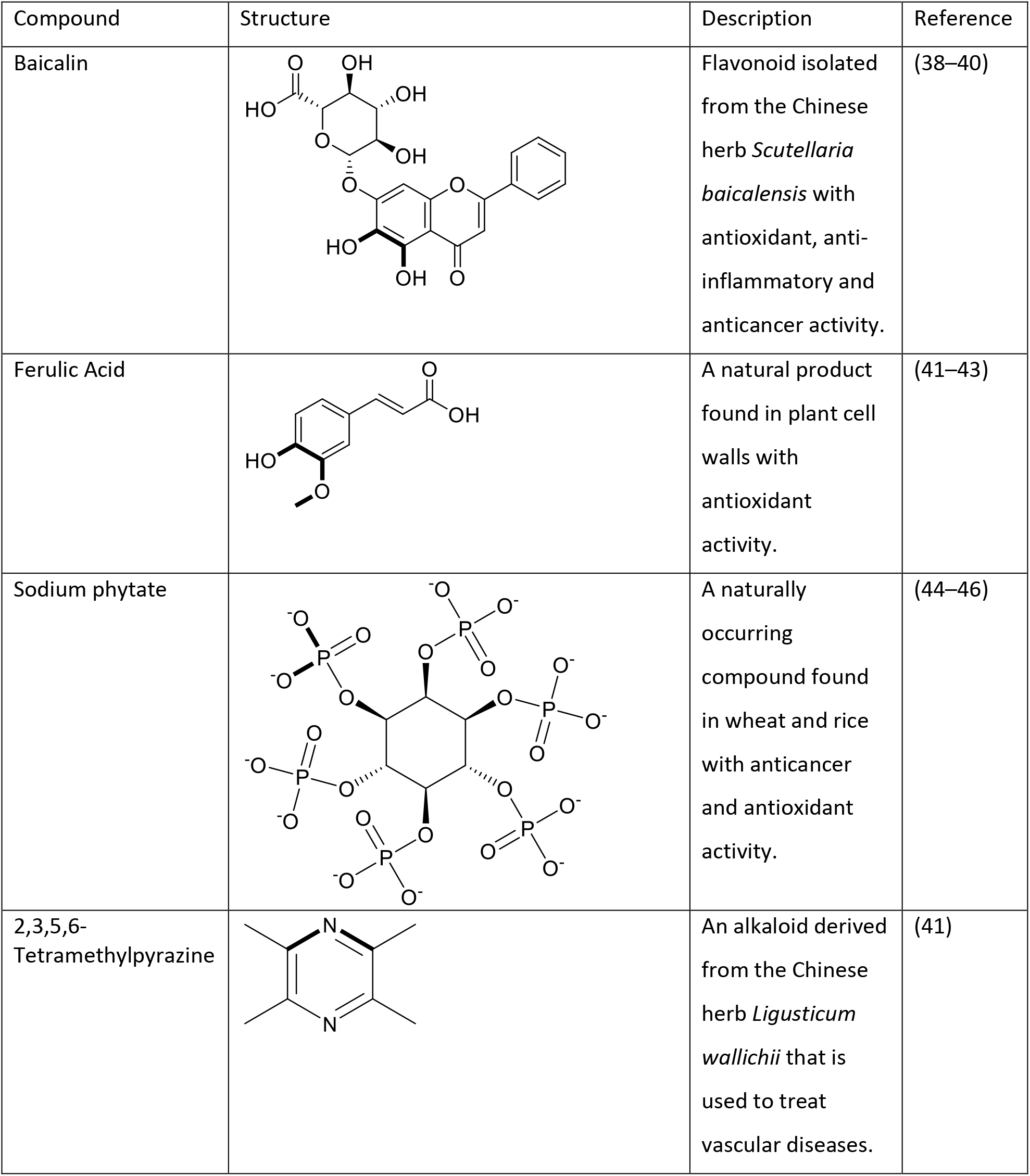
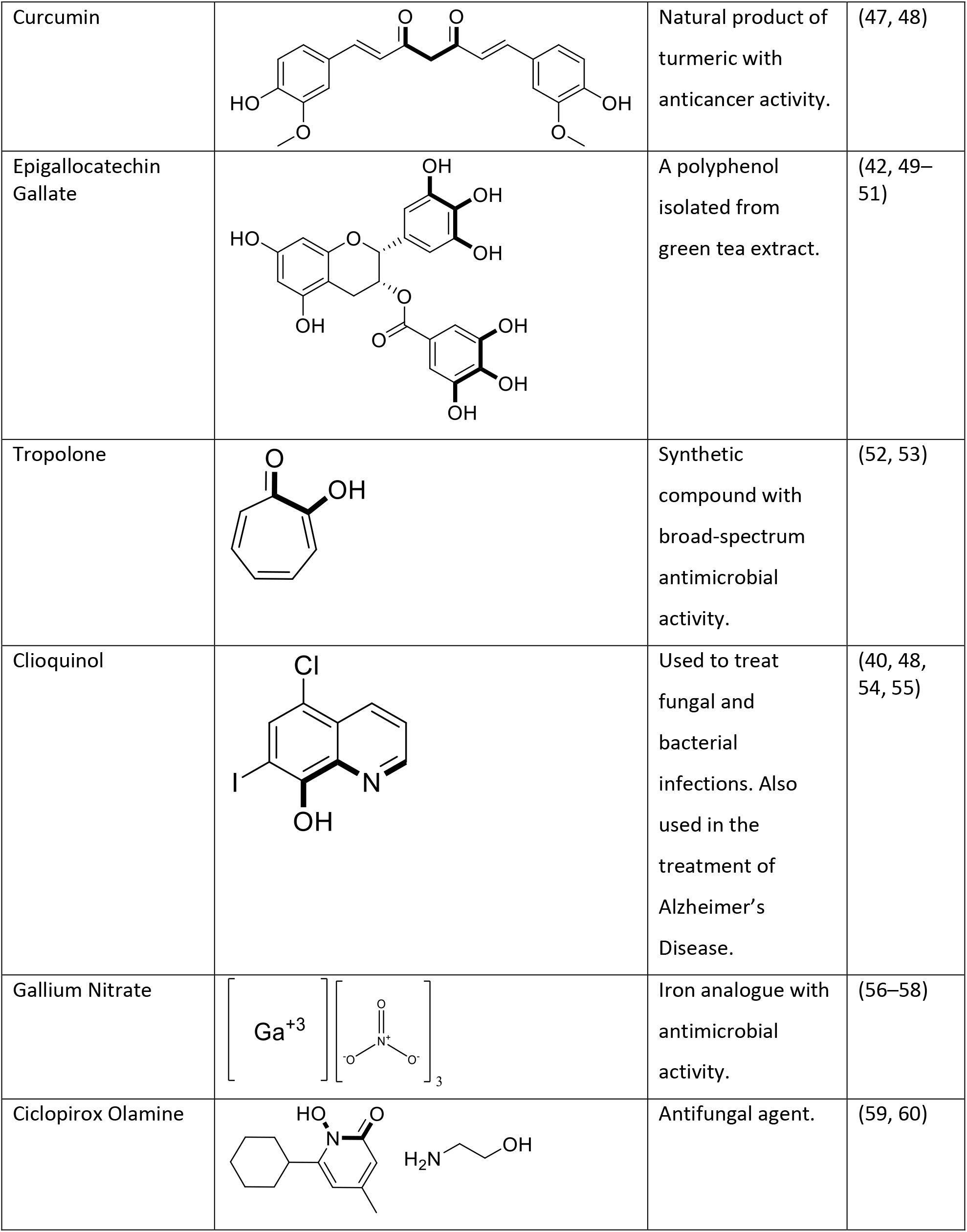
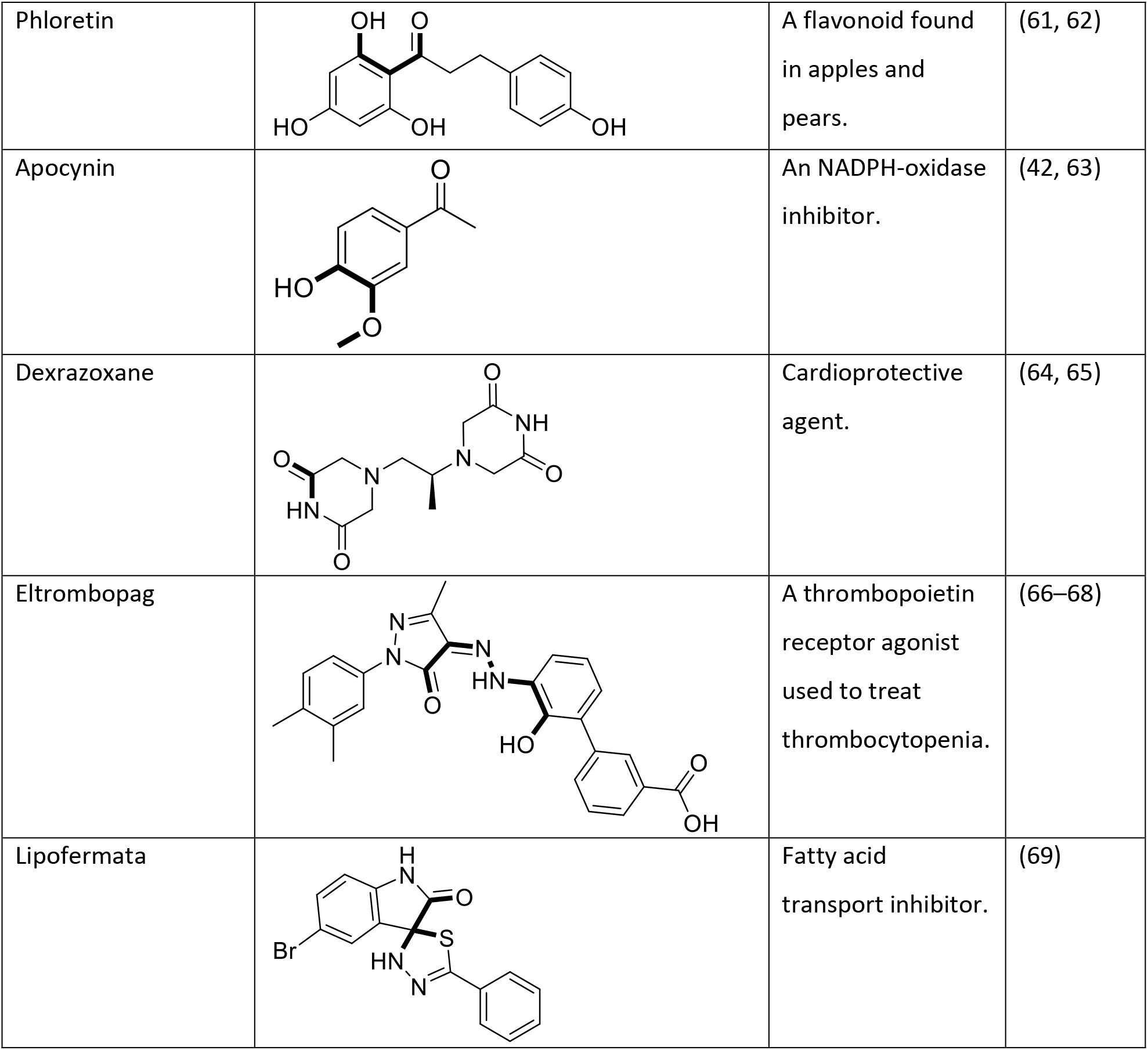
The structures of literature-derived compounds used in this study with potential iron chelation sites bolded.

| Compound | Structure | Description | Reference |
| --- | --- | --- | --- |
| Baicalin |  | Flavonoid isolated from the Chinese herb *Scutellaria baicalensis* with antioxidant, anti-inflammatory and anticancer activity. | (38–40) |
| Ferulic Acid |  | A natural product found in plant cell walls with antioxidant activity. | (41–43) |
| Sodium phytate |  | A naturally occurring compound found in wheat and rice with anticancer and antioxidant activity. | (44–46) |
| 2,3,5,6-Tetramethylpyrazine |  | An alkaloid derived from the Chinese herb *Ligusticum wallichii* that is used to treat vascular diseases. | (41) |
| Curcumin |  | Natural product of turmeric with anticancer activity. | (47, 48) |
| Epigallocatechin Gallate |  | A polyphenol isolated from green tea extract. | (42, 49–51) |
| Tropolone |  | Synthetic compound with broad-spectrum antimicrobial activity. | (52, 53) |
| Clioquinol |  | Used to treat fungal and bacterial infections. Also used in the treatment of Alzheimer's Disease. | (40, 48, 54, 55) |
| Gallium Nitrate |  | Iron analogue with antimicrobial activity. | (56–58) |
| Ciclopirox Olamine |  | Antifungal agent. | (59, 60) |
| Phloretin |  | A flavonoid found in apples and pears. | (61, 62) |
| Apocynin |  | An NADPH-oxidase inhibitor. | (42, 63) |
| Dexrazoxane |  | Cardioprotective agent. | (64, 65) |
| Eltrombopag |  | A thrombopoietin receptor agonist used to treat thrombocytopenia. | (66–68) |
| Lipofermata |  | Fatty acid transport inhibitor. | (69) |

### Numerous iron chelators synergize with TS

Based on their ability to bind iron, each compound from Table 1, as well as DOXY and CIP, were assessed for synergy with TS using checkerboard assays. DOXY, CO, CLI, TRO, and GN all synergized with TS (Fig. 3), as IC_50_ isobolograms showed that all combinations were below the line of additivity. Combination indices (CIs) were less than 1 (Fig. 3E). Based on the checkerboards, isobolograms, and CI values, CO and CLI demonstrated the most potent synergy with TS while GN had the weakest. Attempts to combine GN with DSX or CO resulted in antagonism, likely due to the chelators binding Ga^3+^ (**Fig. S2**).

**Figure 3.**
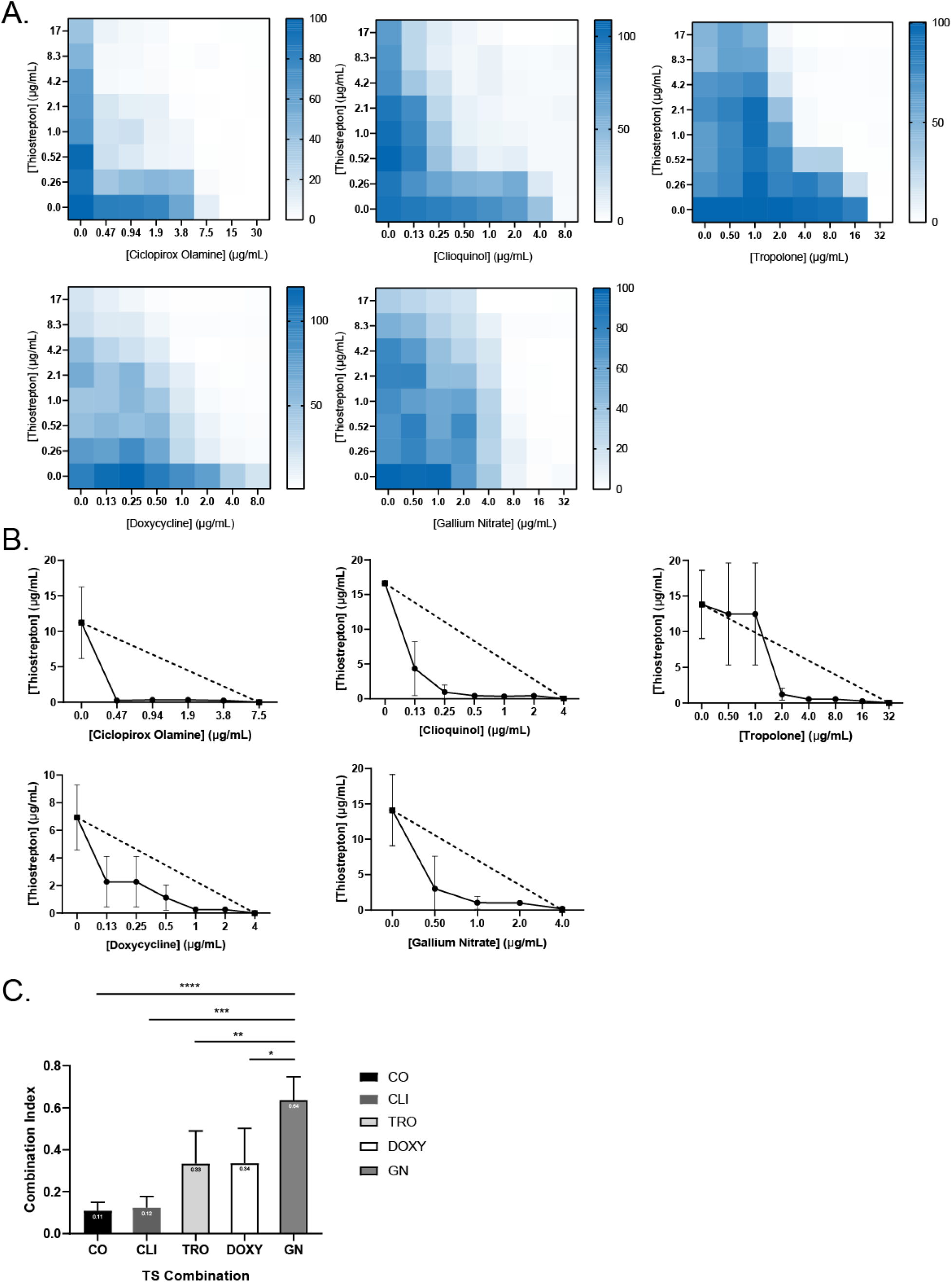
Iron chelators are synergistic with TS against *P. aeruginosa* PA14. **A.** Checkerboards and **B.** IC_50_ isobolograms are shown for each compound that synergize with TS. Dashed lines indicate the line of additivity and solid lines indicate the IC_50_ of TS at each compound concentration. Checkerboards and IC_50_ isobolograms are arranged in the following order: CO, CLI, TRO, DOXY, and GN from left to right, top to bottom. **C.** Combination indices (CI) of each TS combination. CI values are indicated at the top of the bars. All experiments were conducted 3 times. Average values are reported. **** p<0.0001, *** p<0.001, **, p<0.01.

### Each compound potentiates TS activity

We previously showed that iron chelation potentiated the effects of TS, as DSX alone had no anti-*Pseudomonas* activity (22). CLI, TRO, DOXY, and CO can inhibit *P. aeruginosa* growth, suggesting that the innate activity of the compounds could be partly responsible for synergy with TS. Thus, we considered four potential mechanisms of synergy. 1) TS potentiates the activity of each compound through an unknown mechanism. 2) The compound potentiates TS activity by chelating calcium and magnesium and increasing outer membrane permeability or 3) by chelating iron and increasing TS uptake. In all these cases, the synergy is unidirectional. 4) TS and the compound potentiate one another through an unknown mechanism.

Our data suggest that the synergy between TS and each compound is due to their iron chelation capacity rather than membrane permeabilization. First, to determine if DOXY could increase outer membrane permeability, vancomycin (VAN) and DOXY combinations were tested against PA14 alone or in the presence of Ca^2+^, Mg^2+^ or Fe^3+^ (**Fig. S3**). VAN was selected because it is similar to TS in size but unlike TS, its activity is unrelated to iron availability. VAN has a high minimal inhibitory concentration (MIC) against *P. aeruginosa* due to limited uptake across the outer membrane. If a compound increases membrane permeability, we expect synergy with VAN. In our checkerboard assays, no synergy was identified for VAN + DOXY, VAN + CLI, VAN + CO, or VAN + TRO. Further, addition of 100 µM Mg^2+^ or Ca^2+^ had no effect on the checkerboard profiles compared to control. In contrast, addition of 100 µM Fe^3+^ abrogated the inhibitory activity of CLI, TRO, and CO, confirming that iron chelation is a critical part of the mechanism by which those compounds impede growth. Lack of synergy between VAN + DOXY also suggested lack of membrane permeabilization. The addition of 100 µM Mg^2+^ had no effect on the checkerboard whereas the addition of Fe^3+^ and Ca^2+^ had a negligible effect. This is reflective of the relatively weak ability of DOXY to compete for iron in the CAS assay (**Fig. S1**) and of its weak synergy with TS compared to other compounds.

To test the hypothesis that the compounds potentiate TS activity, rather than the other way around, 3D checkerboard assays were performed using PA14. The surface area of each checkerboard was expressed as % of control and graphed against the concentration of the third compound (**Fig. S4**). Individual MIC assays for each compound were performed and the results graphed as % of control on the same y-axis on a log_10_ scale. Significant differences between the two datasets would indicate that the TS + DSX combination potentiates the activity of the test compound. To account for potential antagonism between test compounds and DSX, 2D checkerboard assays were conducted (**Fig. S5**). DSX + TRO and DSX + CO were indifferent. CLI antagonized with DSX at the MIC; however, we could not test concentrations of CLI greater than 8 µg/mL due to its poor solubility. DSX was additive with DOXY. We saw no significant differences between the activity of the chelators alone or in combination with TS and DSX, except for with CLI (Fig. 4ABCD). CLI antagonized DSX at the highest concentration; however, growth was still below 20% of control (Fig. 4C), which we previously established as equivalent to the MIC in the growth medium used for this work (22). When the data were plotted against TS concentration (Fig. 4E), significant differences for the combinations were apparent at 2 and 4 µg/mL TS compared to TS alone, indicating that the compounds and DSX potentiate TS activity. These data suggest that the synergy between the chelators and TS is unidirectional.

**Figure 4.**
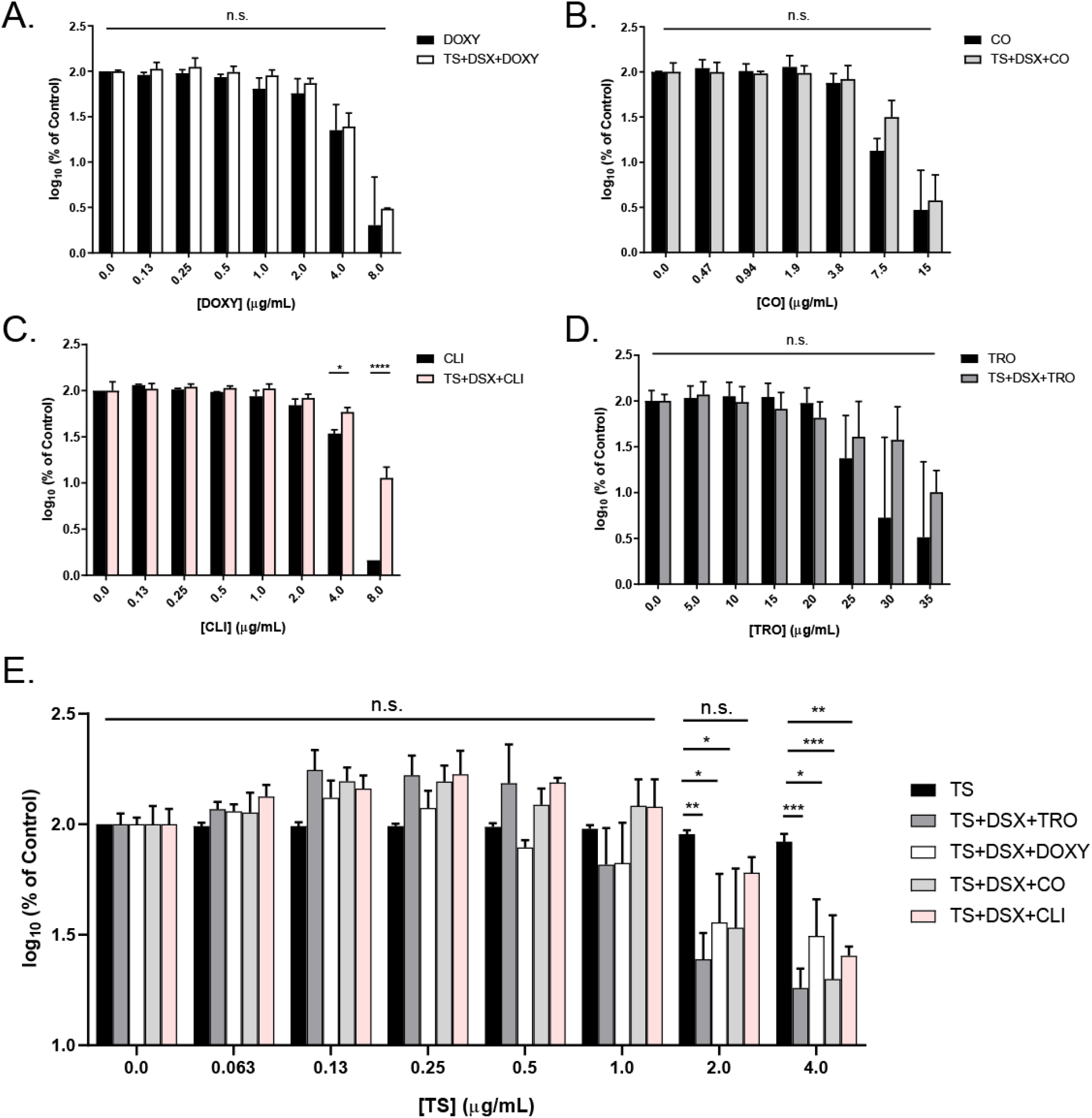
Unidirectional synergy between chelators and TS. Surface areas of 3D checkerboards were plotted against chelator concentration in term of % of control on a log_10_ scale and compared to the activity of each chelator alone. **A.** DOXY. **B.** CO. **C.** CLI. As shown in Figure S4, CLI antagonized with DSX against PA14, thus the triple combination allowed more growth than CLI alone at its highest concentration. However, the triple combination reduces growth below the previously established MIC, 20% of control (21). **D.** TRO. **E.** Surface areas were graphed with respect to increasing TS concentrations and compared to the activity of TS alone. n.s., not significantly different. * p< 0.05, ** p<0.005, *** p<0.0005. The average of at least three biological replicates are shown.

### TS combinations are bactericidal and effective against clinical isolates

TS, CO, CLI, DOXY, and TRO alone were bacteriostatic; however, when combined with TS, the combinations were bactericidal (**Fig. S6**). This improved activity prompted us to test the combinations against clinical isolates. Double (TS + compound) and triple combinations (TS + DSX + compound) were tested against same panels of *P. aeruginosa* and *A. baumannii* clinical isolates we previously assayed for susceptibility to TS + DSX (Fig. 5) (22). GN was omitted due to its weak synergy with TS against PA14 and antagonism with iron chelators (**Fig. S2**). TS and DSX were used at 8.3 µg/mL (5 µM) and 32 µg/mL as before, while the other compounds were added at 1/8^th^ the MIC of PA14, corresponding to DOXY, CO, TRO, and CLI concentrations of 1 µg/mL, 2 µg/mL, 4 µg/mL, and 1 µg/mL, respectively.

**Figure 5.**
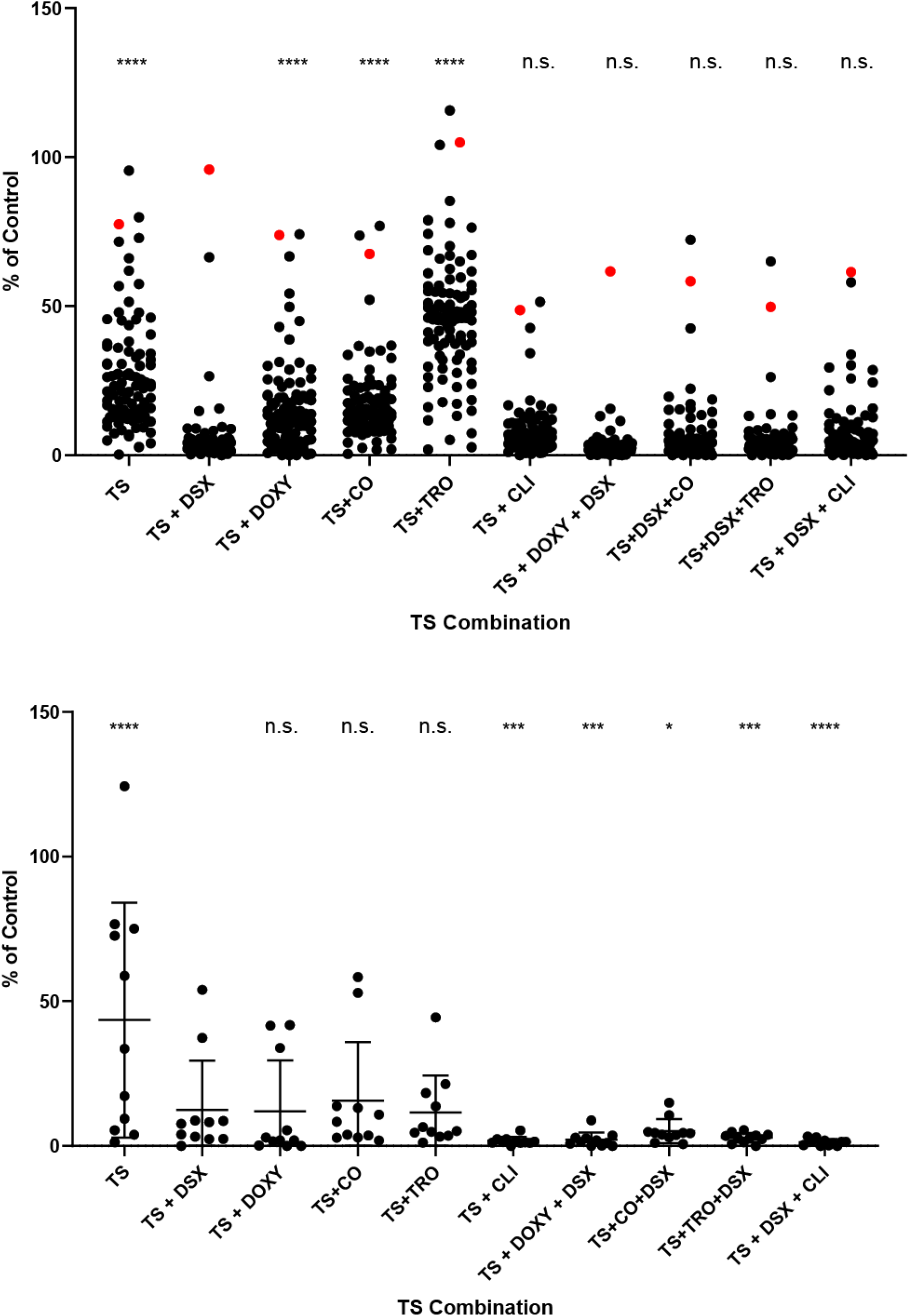
TS combinations inhibit the growth of clinical isolates. Single, double, and triple TS combinations were used to inhibit the growth of **A.** *P. aeruginosa* and **B.** *A. baumannii* clinical isolates. Highly-resistant strain C0379 is highlighted in red. TS and DSX were used at 8.3 µg/mL and 32 µg/mL respectively. The third compound was used at ¼ MIC against PA14 (1 µg/mL DOXY, 2 µg/mL CO, 4 µg/mL TRO, and 1 µg/mL CLI). Horizontal bars show depict the mean % of control growth. Assays were performed at least 3 times. Averaged values are shown. Statistics for TS + DSX versus TS alone or versus other combinations are shown. n.s., not significantly different. *, p<0.05. ***, p<0.0005. ****, p<0.0001.

Of the double combinations, TS + DSX was the most potent against *P. aeruginosa* (Fig. 5A), consistent with our checkerboard assays. Interestingly, TS + DOXY and TS + CO had similar potency despite differences in their CI values (Fig. 3). TS + TRO was the least potent of the double combinations. TS + CLI potency was similar to TS + DSX, and this combination reduced growth of our most resistant clinical isolate, C0379, while TS + DSX did not. TS synergized with CLI to inhibit C0379 although a higher concentration (8 µg/mL) of CLI was required (Fig. 6). Of the triple combinations, TS + DSX + DOXY was the most potent, with only C0379 showing resistance. We previously reported that C0379 has a partial deletion of *fpvB*, encoding a pyoverdine receptor (22). However, triple combinations with higher concentrations of DOXY and CO could inhibit its growth (Fig. 6AC). C0379 growth was also inhibited by TS + CLI or TS + DSX + CLI, if CLI was used at 8 µg/mL. CLI alone did not reduce growth below MIC and there was no antagonism between DSX and CLI with C0379 compared to PA14 (Fig. 6B). C0379 was also less susceptible to TRO compared to PA14 (Fig. 6D).

**Figure 6.**
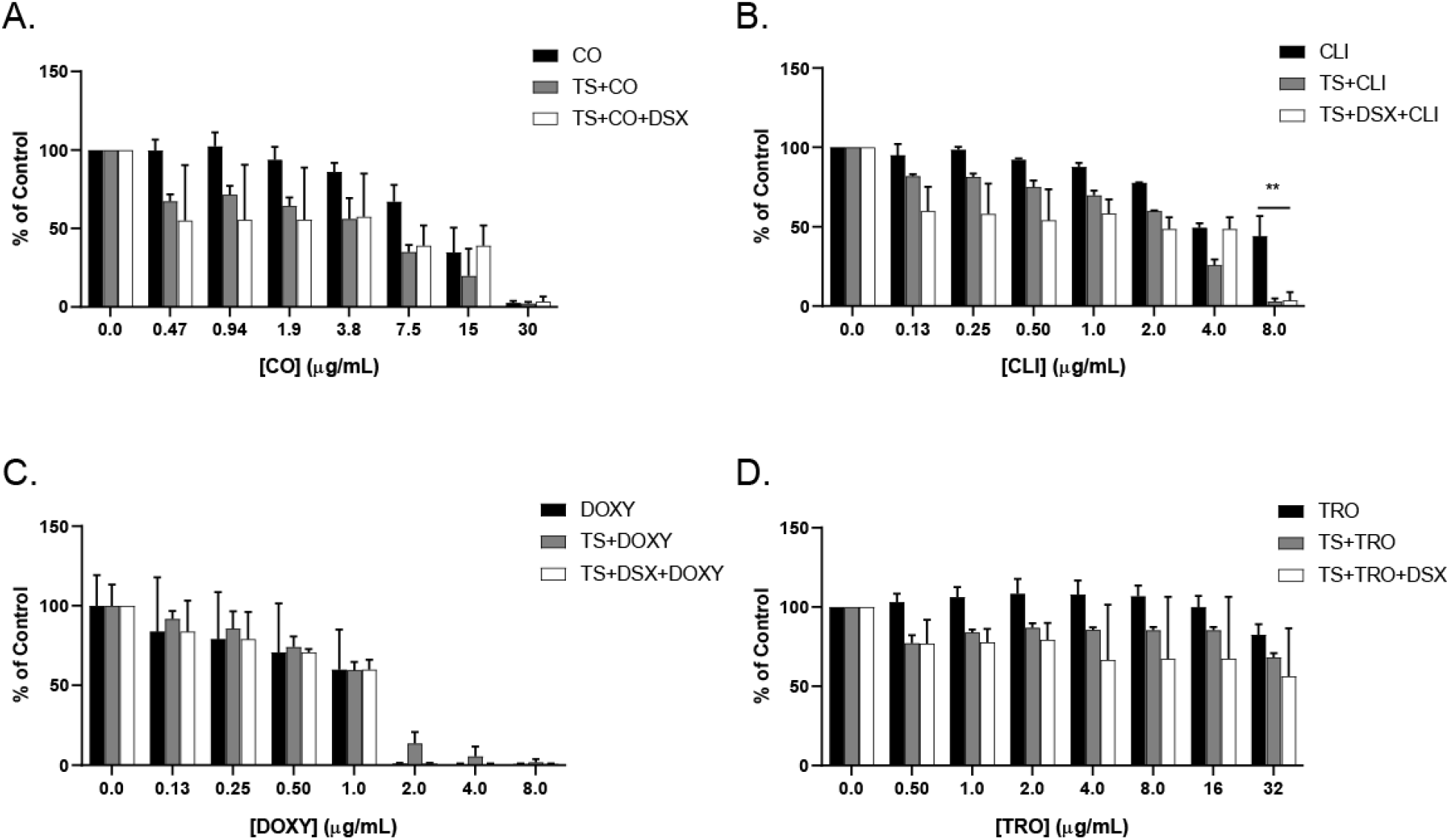
Growth of highly-resistant strain C0379 is inhibited with increased concentrations of CO, CLI, and DOXY. The resistant clinical isolate was challenged with single, double, and triple combinations of **A.** CO, **B.** CLI, **C.** DOXY, and **D.** TRO. Increasing doses of each compound were combined with TS and DSX at 8.3 µg/mL and 32 µg/mL, respectively. MIC assays were conducted at least 3 times and averaged results are shown. **, p<0.001.

For *A. baumannii* isolates, all double combinations were equally effective. TS + CLI was highly potent against *A. baumannii* compared to *P. aeruginosa* when CLI was used at 1 µg/mL (Figs. 5B **and S7**). Strain C0286 was resistant to TS but susceptible to TS + CLI, suggesting inhibition was due to CLI. Conversely, TS + TRO had little activity against *P. aeruginosa* clinical isolates but was effective against *A. baumannii*. The triple combinations inhibited the growth of both species.

## DISCUSSION

A diverse repertoire of iron-uptake mechanisms allows *P. aeruginosa* to proliferate under iron-limited conditions, similar to those encountered during infection of a host. Thus, repurposing iron chelators as antibiotic adjuvants may increase expression of iron-uptake pathways that can then be exploited to deliver antibacterial compounds. Both *P. aeruginosa* and *A. baumannii* express pyoverdine receptors FpvA and FpvB, which are highly upregulated under iron-limited conditions (6, 7). TS hijacks these pyoverdine receptors to enter the cell, as mutants lacking both receptors are resistant. The combination of TS + DSX inhibited the growth of most clinical isolates (22). Here we identified additional iron-chelating compounds that synergize with TS to inhibit growth of the few clinical isolates that were resistant to TS + DSX.

Antimicrobial-iron chelator combinations have been explored for treatment of both bacterial and fungal infections. A combination of DSX and tobramycin inhibited *P. aeruginosa* biofilm formation on CF airway cells (27), while chelation of iron by DOXY potentiated the activity of fluconazole against *Candida albicans* (28). For *P. aeruginosa*, iron restriction has the added benefit of increasing twitching motility and reducing biofilm formation, leaving cells more susceptible to antibiotic treatment (29).

Here we identified multiple compounds that synergize with TS against *P. aeruginosa* and *A. baumannii* clinical isolates, due to their ability to chelate iron. Iron-binding capacity was demonstrated by monitoring visual color changes when complexed with Fe^3+^, CAS agar decolorization, and via spectrophotometric assays. The CAS assay, which is used to detect siderophore production, not only indicates whether a compound can bind iron, but also if it has a stronger affinity for the metal than the CAS-HDTMA complex. This allowed us to compare the relative binding affinities of various compounds based on the extent of decolourization. This method is limited by compound solubility, as seen with CLI (**Fig. S1)**.

None of the natural phytochelators from plants that we tested – including baicalin, ferulic acid, sodium phytate, 2,3,5,6-tetrametylpyrazine, curcumin, epigallocatechin gallate, and phloretin (Table 1) – synergized with TS. *P. aeruginosa* can act as a plant pathogen and may have evolved to outcompete or even take up phytochelators (30–33). The compounds that synergized with TS are all synthetic and the extent of synergy correlated with their ability to strip iron from CAS-Fe^3+^-HDTMA complexes (Fig. 3 and **Fig. S1**). Iron chelators compete with siderophores and reduce iron availability, resulting in increased pyoverdine receptor expression and susceptibility to TS (34). Weaker chelators such as DOXY and CIP showed little or no synergy with TS whereas strong chelators like CO and TRO exhibited greater synergy.

The GN data demonstrate that synergy with TS can occur via routes other than iron chelation. Ga^3+^ represses pyoverdine production and forms complexes with pyoverdine that prevents iron binding (35, 36). TS activity could be weakly potentiated because of reduced competition for pyoverdine receptors if siderophore production decreases upon GN treatment. These data show that disrupting iron acquisition may be another avenue for novel TS combinations. GN in triple combinations with TS + chelator has limited utility because iron chelators bind Ga^3+^ (**Fig. S2**). However, one study showed that LK11, a compound that inhibits pyoverdine function directly, sensitized cells to CO to the same extent as a pyoverdine null mutant (37). Our previous work showed that a PA14 *pvdA* transposon mutant was more susceptible to TS compared to the wild type, which suggests that pyoverdine biosynthesis inhibitors could be useful TS adjuvants (22).

In summary, TS synergizes with iron-chelating compounds of diverse structure that were not primarily intended as antibacterial compounds. Although the mechanisms of action for some of these molecules are not fully understood, they may reveal new targets for antibiotic therapy. In addition, TS combinations demonstrated bactericidal activity while chelator compounds alone were bacteriostatic. The new combinations were effective against clinical isolates resistant to TS+DSX. Our data suggests that these compounds have dual roles – as antibacterial agents and TS adjuvants. Iron restriction mimics many *in vivo* conditions, as host proteins sequester free iron in an attempt to starve bacteria and exert antibacterial activity. Screening for antibiotic activity under similar conditions is an important strategy for development of new treatments for the most dangerous pathogens.

## ACKNOWLEDGEMENTS

We thank Gerry Wright for access to strains from the Wright Clinical Collection and Jakob Magolan for synthesis of DSX. This work was funded by grants to LLB from the Natural Sciences and Engineering Research Council (NSERC) RGPIN-2016-06521 and the Ontario Research Fund RE07-048. DC holds an NSERC Canadian Graduate Scholarship – Master’s Award. IG held a Summer Studentship from Cystic Fibrosis Canada.

## METHODS

### Bacterial strains, compounds, and culture conditions

*P. aeruginosa* PA14 was used for checkerboard assays as previously described (22). All clinical isolates were from the Wright Clinical Collection as described previously. Bacteria overnight cultures were grown in lysogeny broth (LB) and subcultured in 10:90 medium (10% LB, 90% phosphate buffered saline (PBS)). All growth assays were done using 10:90 and grown for 16h at 37°C and 200 rpm. Compounds from Table 1 were from AK Scientific, Sigma, and Cayman Chemicals. TS and DSX were from Cayman Chemicals and AK Scientific respectively. Compounds were stored at −20°C. Stock solutions were stored at −20°C until use except for the tetracyclines, which were made fresh due to precipitation at −20°C.

### Absorption spectra assays for iron chelation

Compounds were arrayed in Nunc 96 microwell plates. Vehicle controls contained Milli-Q H_2_O with 1:75 dilution of each compound at a final concentration of 300μM. The experimental set-up contained the same components as the vehicle control, with the addition of 300μM FeCl3. The final volume in each well was 150μL. The plate was incubated at room temperature for one hour and absorption spectra from 300 nm to 700 nm was read in 2nm increments (Multiskan Go - Thermo Fisher Scientific).

### CAS assay

CAS agar plates were prepared as described previously (22). Compounds were standardized to 2 mg/mL and 10 µL of each was spotted onto the plate. Plates were incubated at room temperature for 1 h, then photographed. Three replicates were conducted and the image of a representative plate was presented.

### Dose response and checkerboard assays

Dose response and checkerboard assays were conducted as described previously (22). Briefly, overnight cultures were grown in LB for 16 h, 37°C, 200 rpm then subcultured (1:500 dilution) into 10:90 for 6 h. Subcultures were standardized to OD_600_ of 0.10 and diluted 1:500 in fresh 10:90 before use. For the dose response assay, serial dilutions of compounds were added at 75 times the final concentration and diluted with 10:90 with cells to reach the desired final concentration. This was done in triplicate for technical replicates. Vehicle and sterile controls were included. The checkerboard assay was done similarly to the dose response assay but in an 8 × 8 format in a 96-well Nunc plate, with concentration of one drug increasing along the y-axis and the other along the x-axis. Sterility and vehicle controls were included with two columns allocated for each control. At least three biological replicates were repeated for the dose response and checkerboard assays.

### 3D Checkerboard Assays

Three-dimensional checkerboard assays were performed in Nunc 96 microwell plates in an 8 × 8 × 8 matrix format for a total of 512 wells. The first two columns were used for the vehicle controls while the last two columns were allocated to sterility controls, both consisting of 2.7% (v/v) DMSO + 1.3% (v/v) H_2_O for plates with TRO and DOXY and 4% (v/v) DMSO for plates with CLI and CO. Serial dilutions of TS were added along the y-axis of each plate starting at 0 μg/mL, with the highest final concentration being 4 μg/mL. Serial dilutions of DSX were added along the x-axis of each plate, from 0 μg/mL to the highest final concentration of 8 μg/mL. Serial dilutions of compound were added with an increasing concentration in each plate up to a final concentration of 35 μg/mL (TRO), 8 μg/mL (DOXY), 30 μg/mL (CO), and 8 μg/mL (CLI) in the last plate. Each well contained 144 μL of 10:90 inoculated with PA14, except for the sterility control columns which contained 10:90 only. The final volume in each plate was 150 μL. The plates were sealed with parafilm and incubated at 37°C for 16 h, shaking at 200 rpm. The OD_600_ of the plates was read (Multiskan Go - Thermo Fisher Scientific). Each experiment was repeated at least three times. Checkerboards were analyzed in Excel. Representative plots at ¼ MIC were made using MATLAB. Surface areas were averaged, expressed in % of control, and plotted against each compound concentration (Prism, Graphpad).

### Clinical Isolate Testing

Isolates from the Wright Clinical Collection were grown and tested as described previously (22). Briefly, clinical isolates were inoculated from glycerol stocks stored at −80°C into Nunc 96-well plates and grown overnight at 37°C, for 16 h with shaking in LB (200 rpm). Overnights were subcultured (1:25) into fresh 10:90 medium and grown for 2 h under the same growth conditions. Subcultures were diluted 1:75 in fresh 10:90. Compounds were diluted 1:75 to obtain the final concentration. DOXY and CLI were added at a final concentration of 1 µg/mL, CO was used at 2 µg/mL, TRO was used at 4 µg/mL, TS was used at 8.3 µg/mL, and DSX was used at 32 µg/mL. Vehicle and sterility controls were included. Plates were incubated overnight with the same conditions. The OD_600_ was read (Multiscan Go – Thermo Fisher Scientific), analyzed using Excel, and the data plotted using Prism (GraphPad).

